# Soil microarthropod community assembly at the micro spatial scale - A microcosm manipulation study

**DOI:** 10.1101/2024.10.14.618259

**Authors:** T. Dirilgen, T. Bolger

**Affiliations:** School of Biology & Environmental Science, University College Dublin, Belfield, Dublin 4, Ireland; Department of Biology, Maynooth University, Maynooth, Co. Kildare, Ireland; Earth Institute, University College Dublin, Belfield, Dublin 4, Ireland

**Keywords:** Community dynamics, dispersal, soil biodiversity, Collembola, mites

## Abstract

Our understanding of soil microarthropod (Acari and Collembola) community assembly and dynamics is somewhat limited compared to aboveground communities. Understanding the processes involved in assembly and the spatial scales at which they occur would help answer the age old question of how so many species and individuals can coexist in soil. We use a microcosm experiment using intact soil cores to explore the processes of selection and dispersal taking place at the micro-spatial scale. We do this by manipulating available pore space and population density, which allows us to indirectly investigate the role of dispersal and biotic interactions in shaping microarthropod community dynamics. Results suggest that there are processes limiting abundance and that communities are sometimes held at abundances below those which the environment could accommodate by abiotic factors. Food and space did not appear to drive the observed patterns; however, findings suggest that abiotic factors may influence dynamics in the field.

## 1 Introduction

The below-ground portion of the terrestrial ecosystem is amongst the most biologically diverse environments and harbour approximately 25% of global biodiversity in terms of species (Decaëns 2010, Coleman and Wall 2015 (FAO et al., 2020). Many of these below-ground organisms are widely distributed and occur in large numbers making them integral components of many ecosystem functions (Trap et al. 2016; George et al. 2019).

Although ecologically and functionally important, soil organisms have had a negligible influence on the development of contemporary ecological theory (Wardle and Giller, 1996; Eijsackers, 2001; Thakur et al. 2020). There is growing evidence that soil communities are structured at multiple spatial scales by a number of interplaying abiotic and biotic factors, with the relative roles of these factors changing across scales (Caruso, Schaefer, Monson, & Keith, 2019; Davison et al., 2015; Ettema & Wardle, 2002; Vályi, et al., 2016). Because these “acts” (interactions) are played out on various spatial and temporal scales in what Hutchinson (1965) called the ‘ecological theatre’, in order to interpret the drama, one must view each component at an appropriate scale (Ecology et al. 2007).

The framework proposed by Vellend (2010) suggests that four processes; 1. selection (i.e. deterministic fitness difference between individuals of different species e.g. environmental filtering), 2. drift (i.e. random changes in species relative abundances), 3. speciation (i.e. the creation of new species) and 4. dispersal (i.e. the movement of organisms across space), interact to determine community dynamics across these different spatial scales. A recent review by Thakur et al. (2020) shows that these processes have been investigated to varying extents across the spectrum of soil taxa, with some processes and some organisms being studied more than others. It is only recently, that Vellend’s community assembly framework is used in the context of below-ground organisms, specifically for microbial communities (see Cordovez et al 2019 review).

There has been relatively little work done on soil microarthropod community assembly dynamics (see nice review outlining gaps by Potapov et al. (2020) and gaps highlighted by Thakur et al. (2020), show that (i) interaction-based processes (i.e. selection) and, (ii) soil mesofauna (microarthropods) are the least represented group. Microarthropods (Collembola and Acari) are often the most abundant component of the soil fauna (Bardgett, 2005) with densities as high as 200,000 m^-2^ in agricultural grassland (Bolger and Curry, 1984). They are part of the complex belowground subsystem which is important in the processes of decomposition and nutrient mobilisation and thus contribute to both provisioning and regulating ecosystem services (Decaëns et al., 2006; Lavelle et al., 2006; Mulder et al., 2011). Despite their observed high densities, these numbers must somehow be kept within certain ‘predetermined’ bounds (c.f. Power, 1992 review) and perhaps be returned to some predetermined mean density following disturbance (White, 1978). However, we know little about whether soil fauna are ever at their limit. Factors such as the soil environment, disturbance, resource availability, biotic and trophic interactions are known to affect the abundance of soil fauna. As both spatial and temporal fluctuations in the size of natural populations are common and well documented phenomena, determining the forces involved in regulation are difficult to pinpoint. Our understanding of microarthropod (Acari and Collembola) community assembly and dynamics is somewhat limited compared to aboveground and other below-ground communities e.g. microbial community dynamics. In particular there is a lack of testing movement-based processes i.e. dispersal, in intact systems.

One important, yet, overlooked aspect of the soil environment that dictates the process of dispersal, is the physical architecture of the soil. There has been a growing attention on the important role of soil architecture for soil functions (Vogel et al. 2022), trophic interactions (Erktan et al. 2020). Despite recent interest and capacity (thanks to technological advances in X-ray CT) to explore soil architecture and its role in community dynamics, focus to date has been on microorganisms (Pot et al. 2022) . In particular, there has been a shift to investigating processes taking place at the micro-scale for microorganisms (Baveye et al. 2019).

In this study, we begin an explicit investigation into the role of dispersal (limitation) in community dynamics for soil mesofauna. The movement of organisms across space i.e. dispersal can have important consequences in communities. We complement existing field studies e.g. Auclerc et al. (2009) by keeping the physical architecture of the soil intact. We use a microcosm experiment to explore the processes of selection and dispersal taking place at the micro-spatial scale. We do this by manipulating available pore space and population density, which allows us to indirectly investigate the role of dispersal and biotic interactions in shaping microarthropod community dynamics.

The density of microarthropods was increased in intact soil cores, in order to determine whether elevated numbers could be sustained, thus indicating whether the animals are at a ‘limit’ in the given system. The effect of increased microarthropod density on (a) microarthropod population density, (b) community structure and (c) resources such as soil organic matter and microbial biomass was investigated under laboratory microcosm conditions. Additionally the effect of reduction in available pore space is also explored.

## 2 Materials and Methods

### 2.1 Experimental design

Unlike most manipulation experiments that culture and manipulate known species of soil organisms within an artificial soil system e.g. microcosm created using sieved soil or compost – this study is unique in that it 1) uses naturally occurring populations of individuals and 2) keeps the soil physical architecture intact for use as microcosm. Although this is at the cost of mechanistic understanding of the exact interactions taking place, it is a field-realistic proxy of the outcome of processes.

Dispersal (i.e. movement across space) was limited in two ways: first, by placing the microcosm into piping, making movement out of the system not possible, and secondly by manipulating the intact physical soil structure to have reduced amount of pore space. The abundance of individuals in the system was manipulated such that we were able to investigate different levels of population density under dispersal limitation across time. The role of selection was directly investigated by increasing the density of individuals in a defined unit of space. Food resources available within the environment were measured as an indirect measure of the process of selection e.g. bacteria and fungi as food source.

The study used a fully orthogonal four-factor experimental design where each factor had two levels. The four factors were baseline, density, time and porosity. In order to create different starting baselines, sampling of intact soil cores for use as microcosms took place during two different seasons. This allowed us to test our hypotheses by having two different starting population densities and assemblage structures Baseline 1 and Baseline 2 with density D1a and D1b respectively). The densities were either the ambient field density from a soil core (Control, D1) or an elevated density which was created by adding animals extracted live from seven additional soil cores (High, D8). The abundance of microarthropods within the experimental microcosms were assessed at two time points, at the start (Time 0) and after 120 days (four months, Time 4). Two levels of soil porosity were used, that which occurred in the field and a reduced porosity achieved by compressing the soil cores. Five replicate soil cores were examined for each treatment on each sampling occasion. Soil porosity/pore volume was used to quantify available pore space (*n*=2). Food as resource in the environment was assessed by measuring microbial biomass and composition, and soil organic matter (SOM) (*n*=3).

The experiment had a total of 116 microcosms. Sixty-four microcosms from Baseline a (2 densities X 2 times X 2 porosities X 8 replicates (five for microarthropods and three for food resource). In Baseline b, microbial biomass and SOM were only assessed for the control (D1), i.e. 52 microcosms, (2 densities X 2 times X 2 porosities X 5 replicates (for microarthropods), and 1 density X 2 times X 2 porosities X 3 replicates (food resource)).

### 2.2 Sampling

Soil cores were collected from an agricultural grassland, located in Tinahely, Co. Wicklow (Ireland; 52.81092N, -6.55495W). The soil was humic brown earth with a sandy loam texture. The site was previously sampled as part of the Irish Soil Information System project (Creamer, 2014). Sampling was carried out in late spring 2015 (Baseline a) and early spring 2016 (Baseline b) . Intact soil cores (494 in total; Ø 5cm, 5cm height) were taken approximately 30cm apart using a soil corer (300 cores for Baseline a and 194 for Baseline b). Cores were brought to the laboratory (University College Dublin) and stored in an incubator overnight (at 13°C in darkness) prior to use the following day.

### 2.3 Experimental set-up

Grass was clipped from the surface of all the soil cores. The initial field weights of the cores were taken as a baseline measure of moisture/water content (n=10) and water was added to maintain a relatively constant moisture content throughout the experiment. These intact soil cores for use as microcosms were placed in opaque polythene tubing (Ø 5×5 cm) with a gauze bottom (1mm). Each of the experimental units (i.e. microcosms) was placed into an opaque container and incubated at 13°C in darkness for the duration of the experiment. The lids of the containers were closed loosely allowing air to circulate

#### 2.3.1 Limiting dispersal

Intact soil cores (microcosms) were secured within piping, which prevented movement in and out of the microcosms (from sides and bottom). In addition, some microcosms were mechanically modified to reduce the available pore space available for dispersal. This was achieved by reducing their original heights from 5 cm to 3.5 cm. This was done by applying surface pressure uni-axially for 2 minutes to the top of the soil core.

### Measurement of soil pore space

We confirmed that the reduction in the available pore space was achieved by measuring the volume of a range of soil pore sizes. Soil pore volume (Φ_sp_) was calculated for a range of pore size classes where the necessary values were obtained using a centrifugation method (Vero, 2015). In this method the desired/required suction pressures are created using centrifugal force and the volume of water drained between two pressures is divided by the volume of the soil particles; (bulk density (*m*_x_) / particle density (ρ) (where ρ ∼ 2.65) to give the volume of pores for the size class of interest using the equation of Larsen et al. (2004) (Equation 1).

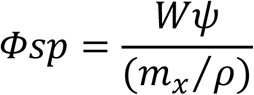

Equation 1 Soil pore volume calculation from Larsen et al. (2004). Where, *Ψ* (cm ^-3^) is the volume of water drained between two tensions, *m_x_* (g) is the mass of oven-dried soil, and *ρ* (g cm^-3^) is the particle density (∼2.65).

The soil microcosms were carefully adjusted to the size required to run in the centrifuge (Ø 4.4 cm) and were saturated with distilled water prior to centrifugation. The soil cores were equilibrated at -10, -56, -91 and -165 kPa suction pressures which roughly correspond to pore sizes of approximately >300, 60-300, 30-60 and 20-30 µm (Nielsen et al., 2008). The centrifuge was a Sigma 6-16KS and the RPM needed to achieve the desired suction pressures were determined based on the unmanipulated soil cores using the Gardner equation (Gardner, 1937). The closest matching values on the centrifuge were chosen, which were; 300, 700, 890, 1,200 RPM respectively.

Pore volume analysis confirmed that available pore space was successfully reduced (Kruskal-Wallis Chi-squared = 11.294, df = 1, p-value < 0.001). The soil cores in which the pore space was mechanically reduced had a significantly lower volume of each of the pore sizes analysed (>300, 60-300, 30-60 and 20-30 µm) (**Fig 1**).

**Figure 1:**
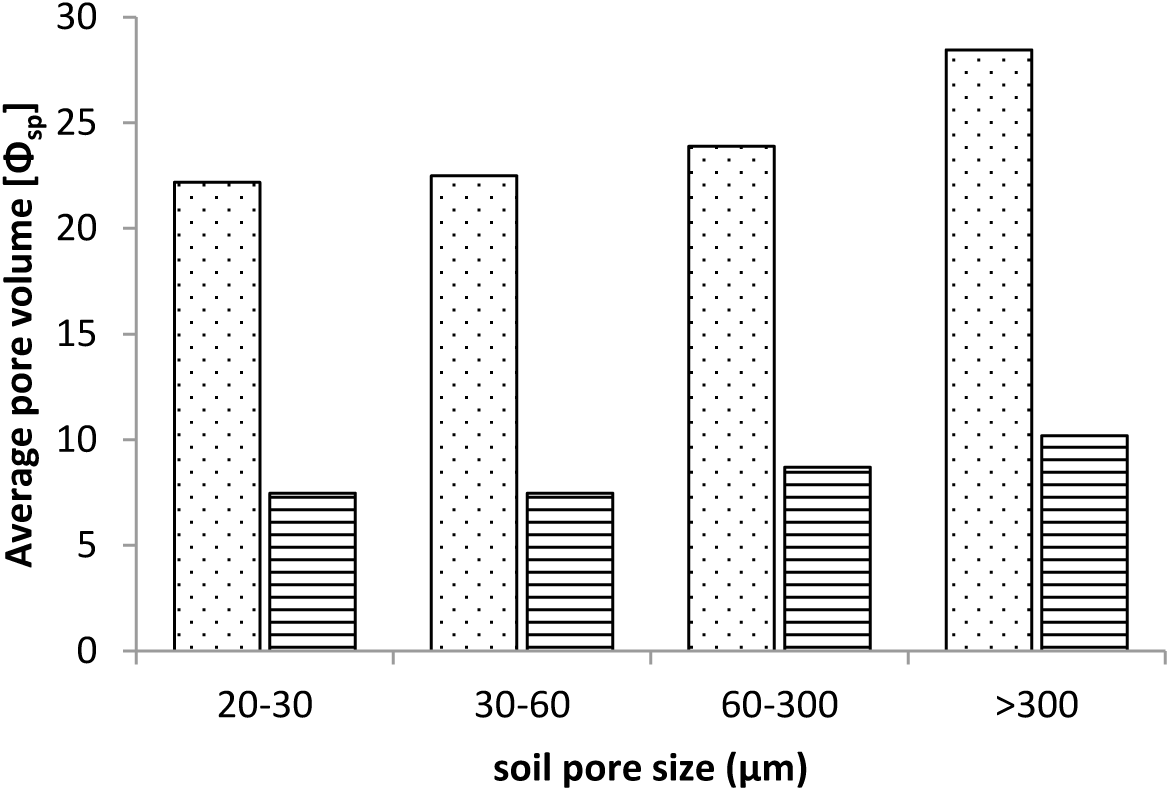
Average pore volume (Φsp) for different sized soil pores 20 – over 300 µm comparing those of control (D1) non-compressed (dots) and compressed (lines) soil cores.(n=2)

**Figure 1:**
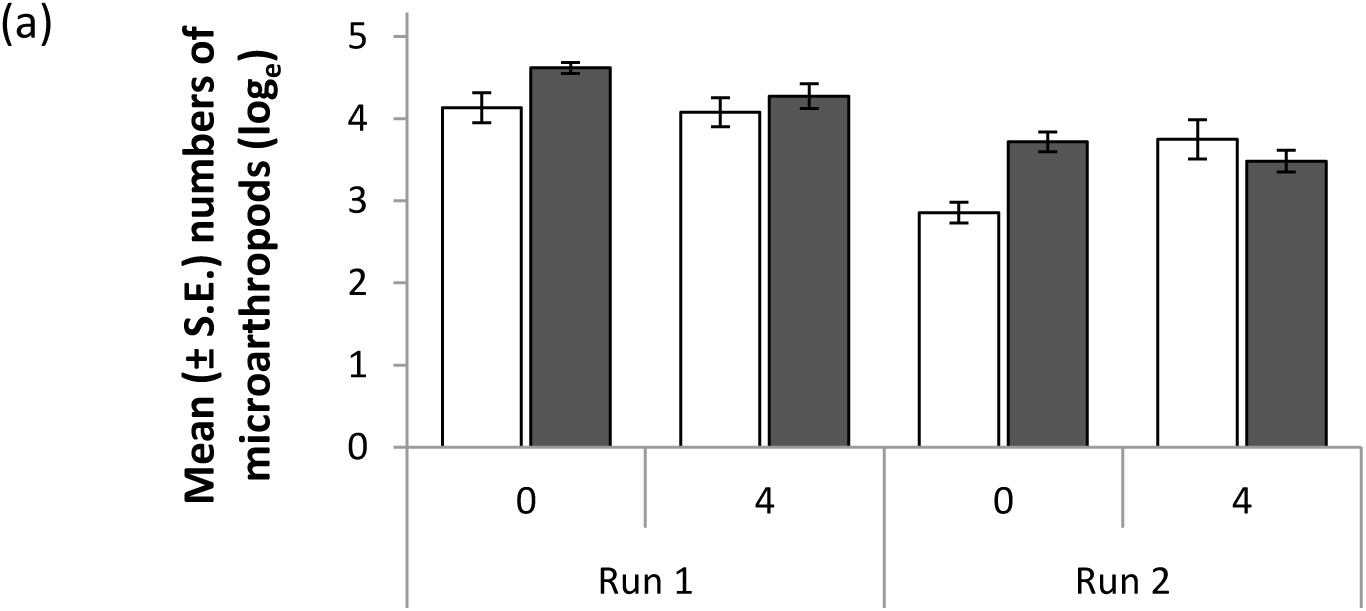
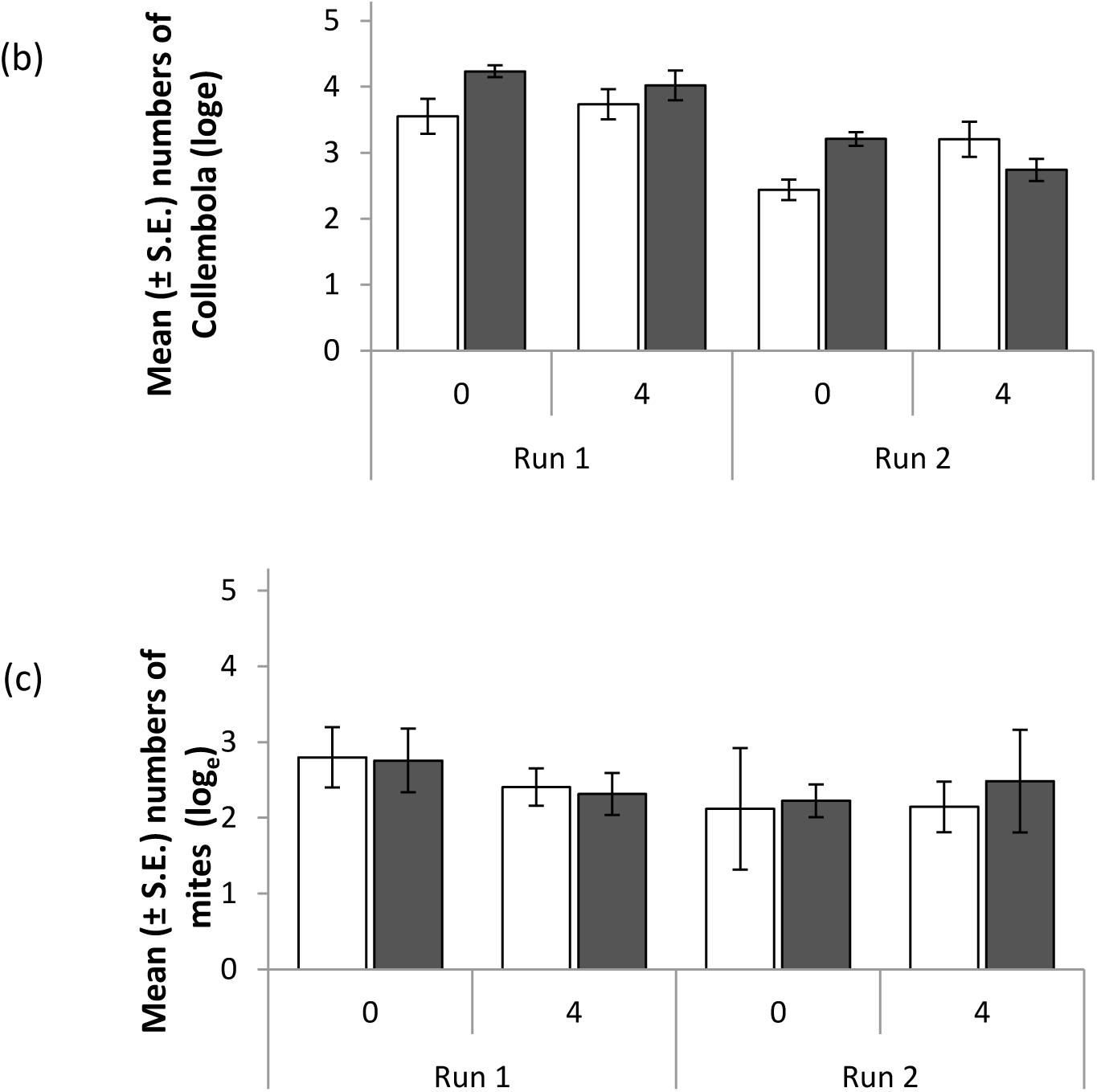
Log_e_ mean (± S.E.) number of (a) microarthropods, (b) Collembola and (c) mites at the beginning (0) and after four months (4) for both baselines of the experiment (Baseline 1 – late spring 2015, Baseline 2- early spring 2016) White bars represent the control treatment and grey bars represent the high density treatment. (*n*=5)

#### 2.3.2 Manipulating and measuring population density and assemblage structure/species composition

In the elevated density treatment (High – D8) microarthropod densities were increased by adding individuals extracted from seven additional cores. The seven cores were extracted together, for live microarthropods, using a High-Gradient MacFadyen extractor into 2 ml of distilled water which was transferred to the top of the experimental microcosm units every second day for 10 days. Two 1ml distilled water additions were used to remove any microarthropods adhering to the sides of the vials into which they were extracted and organisms such as small earthworms were removed. The control microcosms received equivalent water additions (2ml + 1ml + 1ml) at the same times.

At the start (Time 0) and end of the experiment (Time 4) microarthropods were extracted using a Macfadyen High-Gradient extractor into 70% ethanol. Acari (mites) and Collembola (springtails) were slide-mounted in Hoyer’s medium (Krantz, 1978). Where necessary Nesbitt’s fluid was used to clear darker oribatid mites prior to mounting. Specimens were identified to species level where possible using Hopkin (2007) for Collembola and various keys for mites such as those of Weigmann (2006) and Balogh and Balogh (1992) for Oribatida; Karg, (1989; 1993), Evans and Till (1979) and Bhattacharyya (1963) for Mesostigmata; Dindal (1990) for Astigmata; Sig Thor (1933), Gilyarov (1978) and Mahunka (1965) for Prostigmata.

Time 0 measurements were taken 1 week after the animals were added to the microcosms, to allow the introduced individuals time to migrate into the microcosms. Migration into cores was confirmed by visually inspecting microcosms and housing in which they were placed. Time 0 measurements confirmed that the high density treatment (H) contained elevated numbers of Collembola and mite individuals, albeit statistically non significant. An additional experiment confirms that this is an artefact of time 0 measurement date i.e. there is clear evidence that suggests that the number of individuals in the elevated density microcosms decrease within 1 week of being added. Thus, we are confident our analysis and interpretation of Time 0 findings.

We successfully achieved having two differing baselines for the experiment. This was confirmed by the average number of microarthropods in the control microcosms (D1) at the start of the experiment (t1) being significantly different between the two sampling seasons (Tukey’s HSD *p* < 0.05) where the first sampling season (Baseline D1a) had an average of 94.6 individuals and the second (Baseline D1b) 19.2 individuals per microcosm (**Fig 1a**).

### Assemblage structure

One hundred and eighteen microarthropod species were recovered from the microcosms (54 species occurring in both runs, 43 occurring in Baseline a only and 21 occurring in Baseline b only). The microarthropod community composition differed between the first and second baselines (**Fig 2a**) where Monte Carlo tests of the RDA gave a significant effect (pseudo-F=9.4; p < 0.01) which was confirmed by PERMANOVA where there was a significant difference for the main effect of run (F_1,64_ = 5.57; p < 0.01).

**Figure 2:**
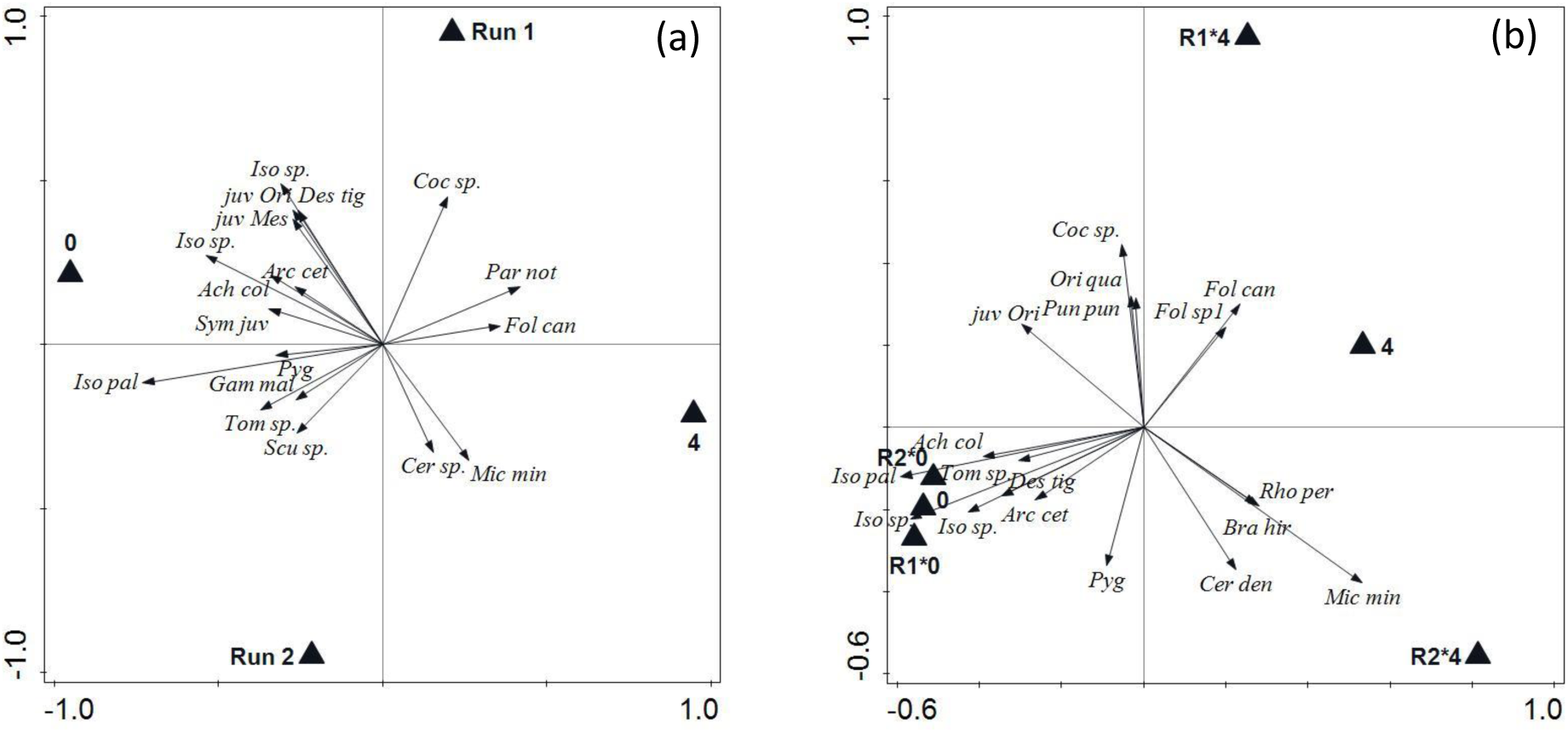
Redundancy analysis (RDA) of (a) microarthropod species abundance and (b) species presence/absence (binary data). Triangles represent factors Time (0 and 4 months), Run (R1 and R2, where Run1 – late spring 2015 and Run2- early spring 2016) and their interaction, ‘*’, if any. .Ordinations show the top 20 species which significantly affected community composition (PERMANOVA; p < 0.05).

#### 2.3.3 Measuring food resource (microbes and organic matter)

Although not explicitly manipulated, we measured micro-organisms (bacteria and fungi) and organic matter in the experimental microcosms, in order to help interpret the findings.

Microcosms were sieved (2 mm) and mixed using the ‘cone and quarter’ technique (Massey et al., 2014). Soil sub-samples to be used for PLFA analysis was stored at -80°C. Microbial biomass carbon (C) was measured using the fumigation extraction method (Brookes et al., 1985; Vance et al., 1987). Sieved soil samples were stored in polyethylene bags and allowed to ‘settle’ of microbial community after disruption by sieving and incubated for 7 days at 21°C prior to fumigation (Bloem et al., 2005). The extracts were analysed for organic carbon using a Shimadzu TOC/TN instrument.

Phospholipid-derived fatty acid (PLFA) profiles were used to compare the composition of the bacterial and fungal assemblages. The ratio of fungal to bacterial PLFAs was used as an index of the relative abundance of these two groups of microorganisms (Leckie, 2005). Samples were thawed at room temperature and immediately freeze dried. The lipid extraction, fractionation and mild alkaline methanolysis were carried out in Coimbra University, Portugal and the fatty acid methyl esters (FAME) were then measured by Gas Chromatography (GC) at Teagasc, Johnstown Castle, Wexford, Ireland. The PLFA biomarkers were deemed to be either, bacteria, fungi, protozoa, eukaryote or plant, according to Frostegård and Bååth (1996), Kaur *et al*. (2005), Frostegård *et al*. (2011) and Francisco *et al*. (2016). Soil organic matter content (%w/w) was measured as loss-on-ignition (LOI) (Allen et al., 1989).

### 2.4 Statistical analyses

Collembola, mite and total microarthropod abundances were analysed using a fully orthogonal four-factor analysis of variance (ANOVA) with fixed factors baseline, density, time and porosity, all of which had two levels. This was done using R version 3.3.2 (R Core Team, 2016) in R studio (RStudio Team, 2015). Four outliers (> 2 S.D. above the mean) were removed and replaced with the average values of the remaining replicates subjected to the same combination of treatments in order to keep a balanced design (following Underwood, 1997). All data required transformation in order to fulfil the assumptions of homogeneity of variance and normality. The Box-Cox method in the “MASS” package (Venables and Ripley, 2002) was used to determine the necessary transformations. Microarthropod and Collembola data were log_e_ transformed and mite data were square root transformed. The “lsmeans” package (Lenth, 2016) was used to make multiple comparisons using the contrast () function. The default Tukey HSD method was used. Kruskal-Wallis rank sum test was used to analyse the pore volume data as they were not normally distributed.

Microarthropod, Collembola, mite species, and PLFA data were Hellinger standardized prior to multivariate analysis. Redundancy Analysis (RDA) was used to detect changes at the species level related to treatment (where treatment was taken to be the explanatory variables and community composition as the response variable). Ordinations were carried out using CANOCO for Windows (version 5) (ter Braak and Šmilauer, 2012). This was followed by a multivariate statistical test to assess significance of differences among treatments. These were evaluated by performing a Permutational Multivariate Analysis of Variance (PERMANOVA) on Hellinger transformed data and Euclidean distance as the distance index.

Unrestricted permutations using the Monte Carlo method were used to assess the significance of the pseudo-F values. PERMANOVA tests were carried out using the adonis () function in the “vegan” package (Oksanen et al., 2017) with 999 permutations. A multivariate analogue of Levene’s test for homogeneity of variances was performed with the betadisper () function using the same package in R.

## 3 Results

### 3.1 Effect of increased population density on

#### 3.1.1 abundance

After the 120 day incubation (Time 4) total number of microarthropods converged, and there were no differences between the two density treatments (Density*Time: F1,64=7.16; p < 0.01). The same was observed for the microcosms with lower starting baseline (D1b), however, the number were less than those from the higher starting baseline. The treatment effect did not differ in microcosms where the available pore space had been reduced (F_1,64_ = 0.16; p = 0.69). We see that this result is driven by Collembola (Figure 1b), as they are the numerically dominant taxon. When only mites are included, their abundance was not significantly effected by any of the four factors. (Fig 1c).

#### 3.1.2 species composition

Differences between the beginning and end of the experiment for both baselines explained most of the observed variation with more species occurring at the beginning (Time 0) of the experiment than at the end (Time 4) where a small group of species dominated (**Fig 2a**) (PERMANOVA, F_1,64_ = 10.16; p < 0.01). For example, *Folsomia candida* Willem, 1902 and *Parisotoma notabilis* Schäffer, 1986 represented 50.87% of individuals at the end and species such as *Microppia minus* Paoli, 1908 occurred in greater abundance at the end of the study than at the beginning. If species presence/absence is used in the RDA the Baseline * Time interactions are significant (**Fig 2b**) and the species composition which was similar at the beginning of both runs diverged markedly by the end (PERMANOVA, F_1,64_ = 4.16; p < 0.01).

The species composition and presence/absence of Collembola were driven by the same variables, i.e. time (**Fig 3 a,b**) (PERMANOVA, F_1,64_ = 13.16; p < 0.01), run (F_1,64_ = 5.24; p < 0.01) and the time*baseline interaction (F_1,64_ = 3.51; p < 0.01). Although the abundance of mites was not significantly related to any of the four factors, the effect on species composition was significant (**Fig 4 a,b**). These ordination patterns were also confirmed by PERMANOVA analyses where time, run and time*run were significant (F_1,64_ = 4.44; p < 0.01, F_1,64_ = 5.62; p < 0.01, and F_1,64_ = 5.51; p < 0.01).

**Figure 3:**
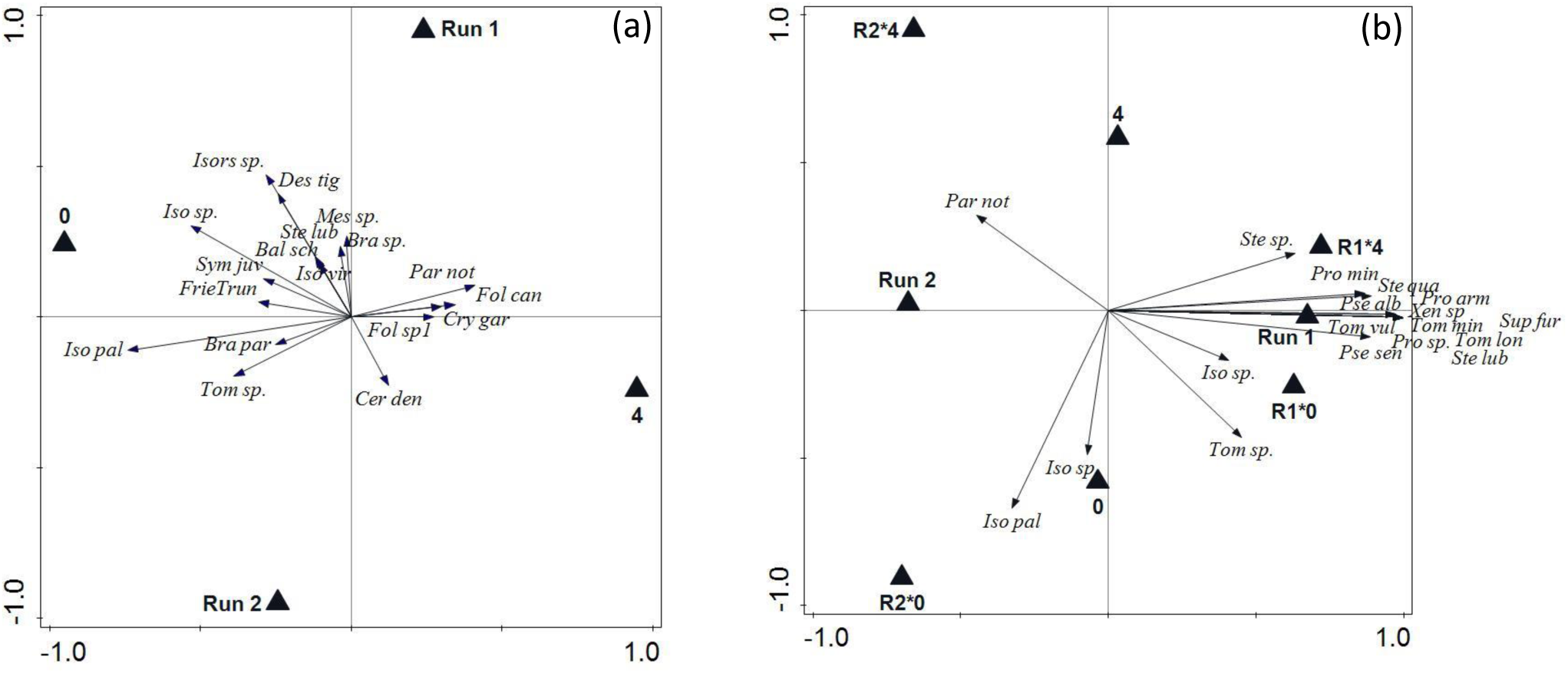
Redundancy analysis (RDA) of (a) Collembola species abundance and (b) species presence/absence (binary data). Triangles represent factors Time (0 and 4 months), Run (R1 and R2, where Run1 – late spring 2015 and Run2- early spring 2016) and their interaction, ‘*’, if any.

**Figure 4:**
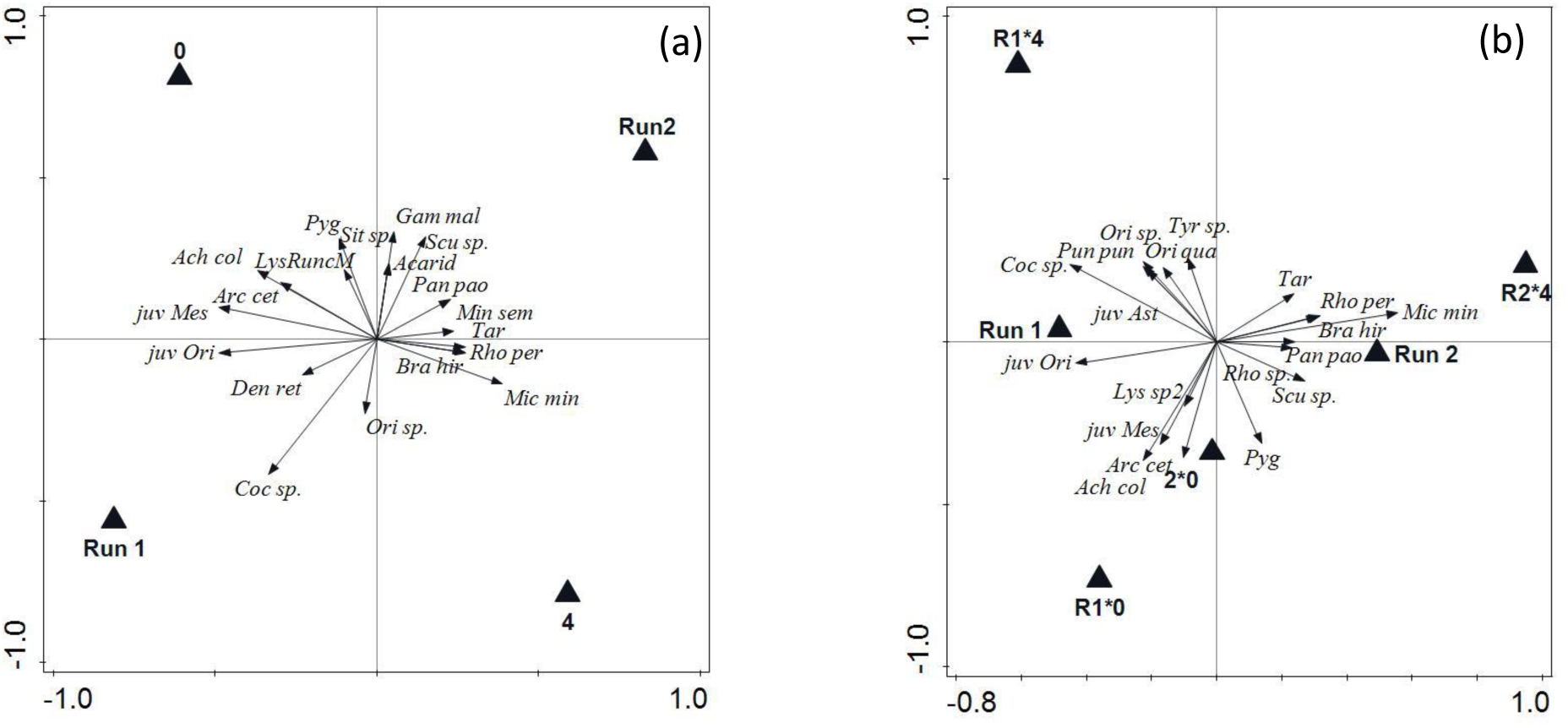
Redundancy analysis (RDA) of (a) mite species abundance and (b) species presence/absence (binary data). Triangles represent factors Time (0 and 4 months)), Run (R1 and R2, where Run1 – late spring 2015 and Run2- early spring 2016) and their interaction, ‘*’, if any.

Additionally, the analyses showed that the interaction between porosity and baseline had a significant effect on microarthropod species composition (F_1,64_ = 1.9; p < 0.05). This effect was not present when Collembola and mites were analysed separately (F_1,64_ = 1.7; p = 0.08 and F_1,64_ = 1.4; p = 0.15). This interaction was not significant for the RDA.

#### 3.1.3 functional diversity

The higher abundance of Collembola in the system meant that fungivores comprised the highest proportion of individuals (∼ 50 %) with predatory individuals making up ∼ 20 % in control samples at the start (**Fig 5**). The abundance of fungivorous microarthropods did not change across time (F_1,78_ = 0.27; p = 0.53) or with reduced pore space (F_1,78_ = 0.44; p = 0.42). Density (F_1,78_ = 4.5; p < 0.04) and Baseline (F_1,78_ = 12.37; p < 0.001) did affect their relative abundance, and they represented a higher relative abundance in the Elevated treatment than the Control (Tukey’s HSD *p* < 0.001). The average abundances of fungivores in both treatments (D1 and D8) were lower in Baseline a compared to Baseline b (Tukey’s HSD *p* < 0.001). On the other hand, numbers of predatory microarthropods remained the same across time for the control while, for both Baselines, numbers decreased in the D8 treatment (Tukey’s HSD *p* < 0.05).

**Figure 5:**
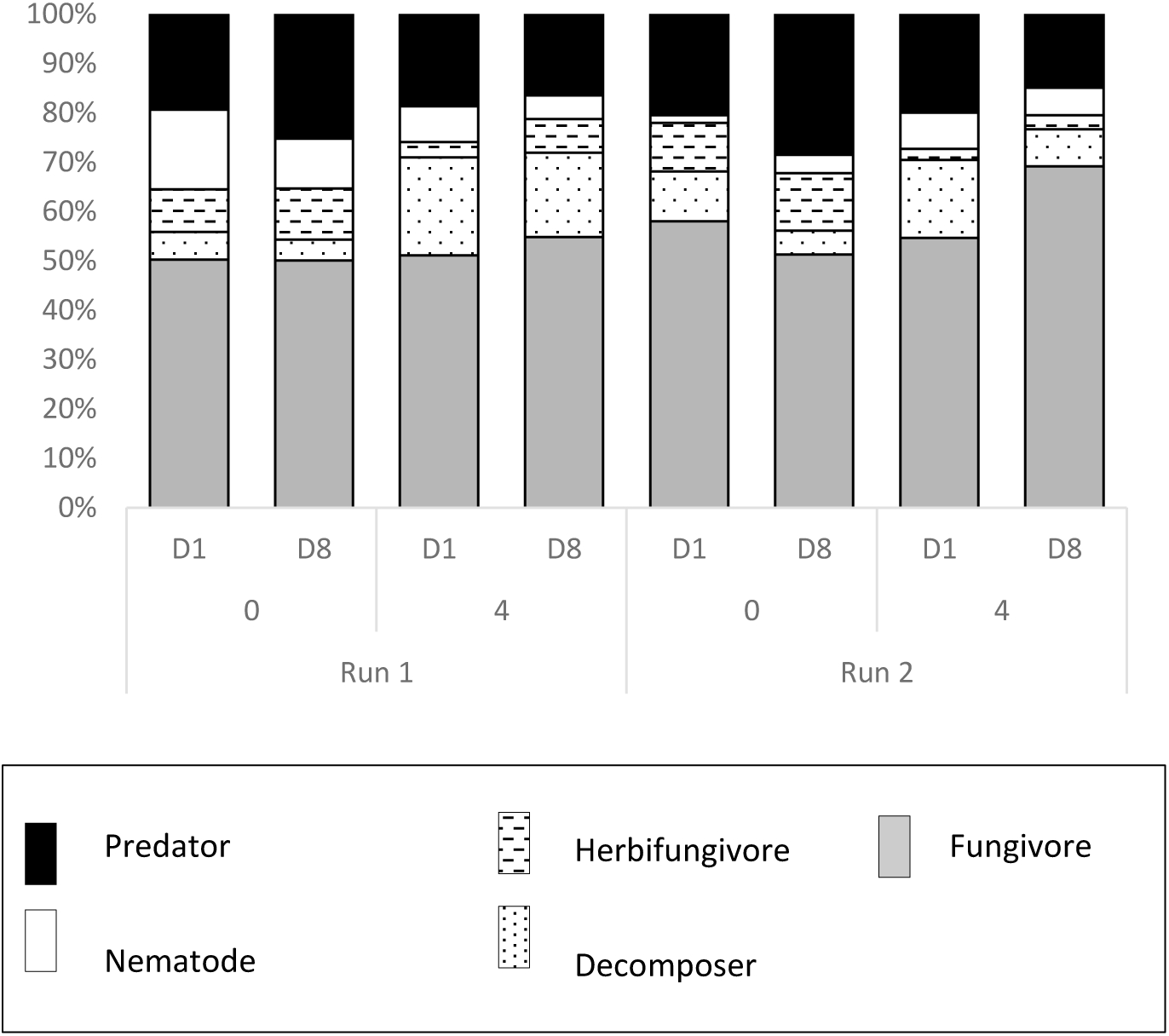
The percentage proportions of microarthropod feeding groups (predator, herbofungivore, fungivore, nematode predator and decomposer) found within the control (D1) and high density (D8) treatment microcosm, at the beginning (0) and after four months (4) for both runs of the experiment (Run 1 – late spring 2015, Run 2-early spring 2016). (n=5)

The nematode predator microarthropods differed between Baselines (F_1,78_ = 12.04; p < 0.001) with the second run containing on average half of the abundance seen in the first run.

The herbivofungivores were composed solely of Symphypleona juveniles and in in the Elevated density treatment, decreased (to zero in some instances) by the end of the 4 month study period (Tukey’s HSD *p* < 0.05). In contrast, the time had an opposite effect on primary decomposer species such as *Folsomia sp.* and *Tomocerus sp.*, where their relative proportion increased at the end (F_1,78_ = 8.86; p < 0.01).

Although reducing available pore space did not affect the overall abundance of microarthropods it did have an effect on some of the feeding groups, i.e. primary decomposers and herbivofungivores, and thus the relative proportion of each group was altered even though the overall abundances did not change significantly.

Effect of reduced soil pore space

#### 3.1.4 Effect on microorganisms (food resource)

In the first run (Baseline a) of the experiment density manipulation did not affect the amount of microbial biomass carbon (F_1,34_ = 0.73; p = 0.49). Microbial biomass in the control (D1) did not differ between Baseline b and 2 (F_1,22_ =3.27; p = 0.08) or between the beginning and end of the incubations (F_1,22_ = 0.59; p = 0.45). Compression did, however, reduce the amount of microbial biomass in the control microcosms (F_1,22_ = 11.37; p < 0.01) (**Fig 6**).

**Figure 6:**
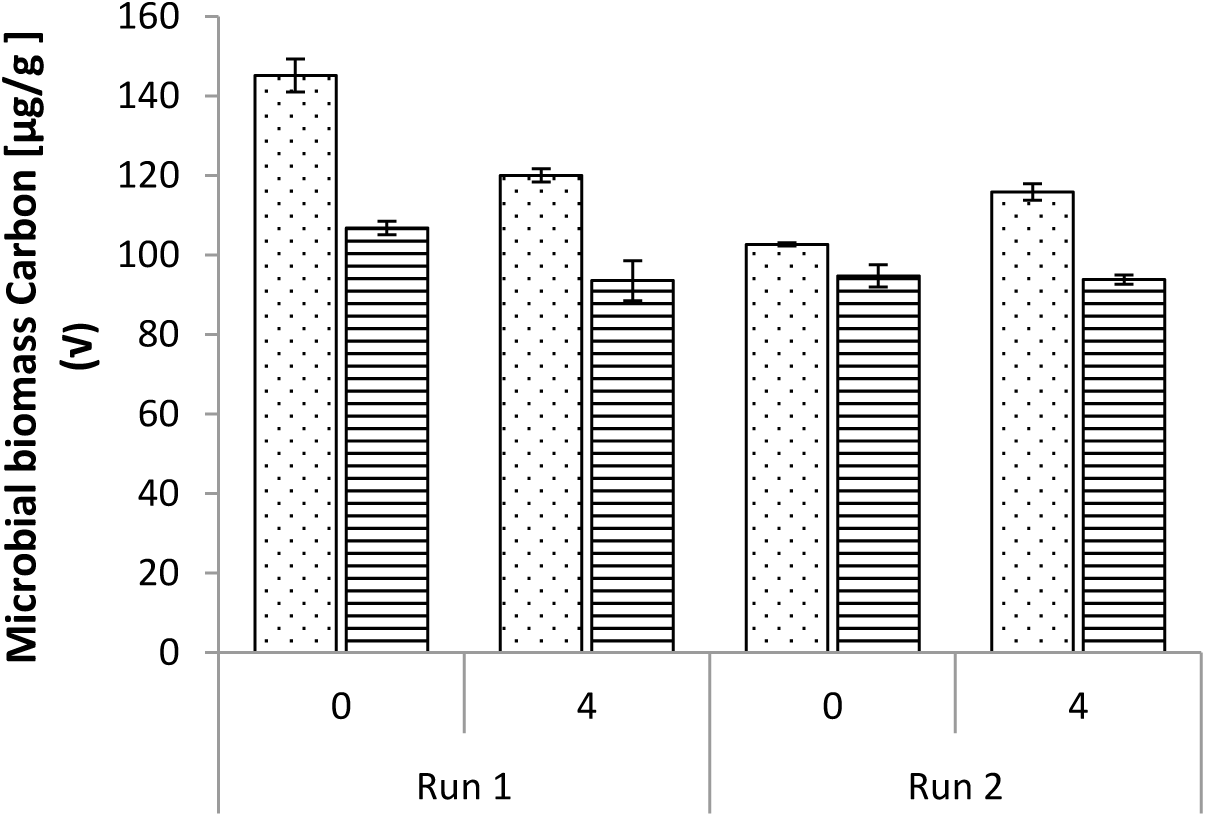
Microbial biomass Carbon (µg/g) for control (D1) non-compressed (dots) and compressed (lines) microcosms at the beginning (0) and after four months (4) for both runs of the experiment (Run 1 – late spring 2015, Run 2-early spring 2016). (n=3)

In Baseline, 1 fungal to bacterial ratio (F:B) was not affected by density, time or compression (F_1,12_ = 0.0003; p = 0.99, F_1,12_ = 1.45; p = 0.25, F_1,12_ = 0.0001; p = 0.99). Similarly, composition of PLFA biomarkers was not related to any of the three factors examined (RDA pseudo F=1.1., p = 0.14). This was also the case when bacterial and fungal biomarkers were analysed separately. However, PERMANOVA of PLFA biomarkers composition did show a significant main effect of density (F_1,12_ = 1.91; p < 0.05). That said, responses to density were the only ones that did not meet the assumption of homogeneity of variance. This can be seen in the RDA graph where the spread of samples is greater in treatment D8 than in D1 (**Fig 7**).

**Figure 7:**
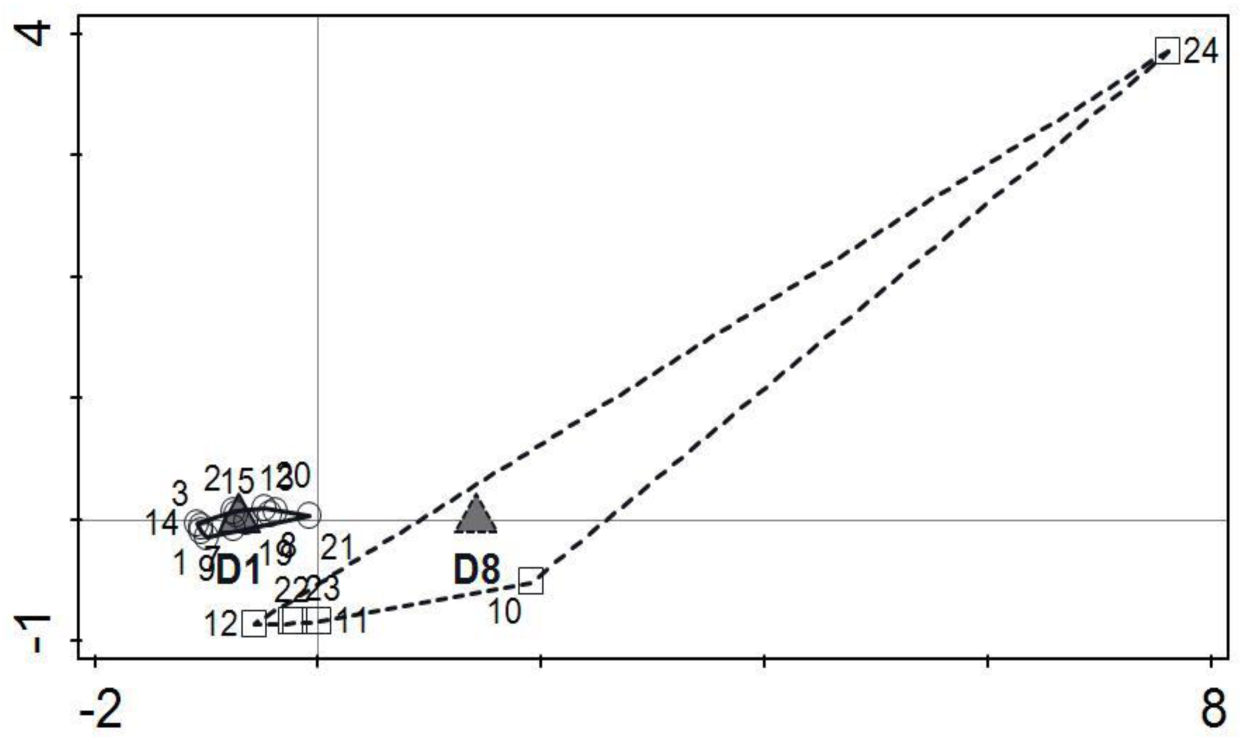
Redundancy analysis (RDA) of PLFA biomarkers showing the spread of samples associated with each density treatment, D1 – control and D8-high density. Both density treatments D1 and D8 are represented by triangles and samples are represented by circles and squares respectively.

## 4 Discussion

Microarthropod abundances in the microcosms appeared to reach a so called ‘limit’ in the first run of the experiment (Run 1) where numbers in the elevated density (D8) returned to that of the control (D1) after a four month incubation period. In contrast, during the second run (Run 2) which had lower starting field densities (∼ 19.2 microarthropod individuals per microcosm), the density in the control treatment increased to those of D8 treatment (∼ 50 per microcosm). This would suggest that this particular agricultural grassland system did have a certain ‘limit’ (∼ 73 individuals in 117.81 cm^-2^) in late Spring, but not operating at its ‘potential limit’ the subsequent year in early Spring. If this is the case and there is a limit to the number of individuals that can be sustained, some factor must be limiting or regulating the given system. It is difficult to decipher which biotic mechanisms might be at play particularly as neither the resources nor predators were specifically manipulated, therefore the response of lower trophic levels to elevated numbers of microarthropods in the system will be used as a proxy to give some indication.

The composition of microarthropods in terms of feeding groups may shed some light on elevated numbers not being sustained (in the case of Run 1) despite there being no (detectable) lack of resources. The increase in microarthropod abundance at the start was primarily due to an increase in fungivorous mites and Collembola, while the abundance of the other functional groups remained the same as the control. The expectation would have been that all feeding groups would have increased in the higher density treatment. This may be an artefact of a combination of two things; firstly, the experimental set-up involved microarthropod additions made every second day over a ten day extraction period which may have been a selective process, i.e. that individuals extracted earlier in the extraction process would have had an increased advantage in entering and ‘settling’ into the microcosms into which they were added. Second, the ten day addition period followed by a seven day ‘rest’ period prior to measuring baseline abundances, may have been ample time for ‘regulating processes’ to be taking place which may explain why we do not see the proportional increase in total microarthropod abundance in the elevated treatment. Indeed, the fact that the densities did not increase eight fold in the D8 treatment further supports the notion that the system may be at some sort of limit. This is supported by a larger increase in the D8 treatment in the second run of the experiment which had smaller starting numbers.

Elevated numbers of fungivores were sustained across the study and there was no indication of a decrease in fungi which would be expected if there were an increase in fungivorous microarthropods. Most microcosm studies investigating fungus-grazer interactions draw a positive correlation between density and grazing pressure/intensity, i.e. the higher the density the higher the grazing pressure. However, these studies generally use single species to examine fungus-grazer interactions in relatively synthetic environments (Bardgett et al., 1993; Hedlund and Öhrn, 2000; Crowther et al., 2012), compared to this microcosm which represents a more natural, multi-species environment. Microbial biomass does not appear to have been over- or under-grazed, which is unlike the findings of similar microcosm studies (such as Chauvat & Wolters 2014). However, this does not necessarily imply that resources were not consumed but rather, it may suggest that the rate/intensity of consumption was such that it provided the opportunity for compensatory growth. Indeed, the ability of fungi to compensate for the effects of grazing has been documented (Mikola and Setälä, 1998), especially when grazing intensity is low (Crowther et al., 2012). Therefore, although microbes do not appear to be top-down controlled, (as is documented in McLean et al., 1996), microarthropods may still be potentially bottom-up (resource) controlled. Indeed, resource manipulation experiments do show food to be a controlling factor (Scheu and Schaefer, 1998; Rantalainen et al., 2004). A potential explanation as to why bottom-up control is not explicitly evident here may be that (i) the use of intact soil cores in the microcosm set-up reflect a more realistic account of what would take place in nature, where microbes possess the ability to compensate for biomass loss, and or, (ii) preferential grazing by microarthropods may have caused a shift in community composition leaving total biomass seemingly unaltered. The latter explanation is not likely as the PLFA did not suggest a change in species composition in the high density treatment. However, the control and high microarthropod density treatments (D8) were not homogeneous in variance for composition of PLFA biomarkers – where there was low variance in the control and high variance in the high density treatment. The fact that variance in microbial species composition increased with increased microarthropod abundance suggests that feeding strategies may have changed as a result of increased competitive pressure, i.e. either more generalist feeders dominate the community or there is a switch to generalist feeding which occurs as abundance increases.

In contrast, although not significantly increased in the treatment with elevated densities, the abundance of predatory microarthropods (i.e. mites of the order Mesostigmata and some Collembola species such as *Friesea truncata*) decreased over time in the higher density (D8) treatment while remaining the same across time for the control. Thus, although the D8 treatment had higher amounts of prey/food (in the form of microbivorous Collembola and mites), this evidently did not sustain or increase predator density. Many studies support the idea that an increase in predators result in the decrease in the density of prey (Lenoir et al., 2007; Schneider and Maraun, 2009). Although predator density was not increased in the elevated microarthrophod density treatment, it is surprising that a larger/higher potential prey density did not sustain or even increase predator density. However, predation rate may not necessarily be influenced by prey population density, where depending on species type predation rate may be positively, intermediately or negatively correlated with prey density (Harris and Usher, 1978). In fact, Schneider et al. (2007) found that increase in the abundance of a predatory gamasid mite was not related to abundance of potential prey. In addition, the microarthropod predators in this study could not be top-down controlled as the larger predators did not occur in the microcosms. Therefore, it may have been expected that predator density would increase as a result of this lack of predator pressure in the control treatment as seen for example in Schneider et al. (2007). However, the strength of predator effects on prey populations may be influenced by intra-guild predation where predators not only attack herbivores but also each other. This ultimately results in a decrease in predator pressure and thus a potential relaxing of top-down control (Finke and Denno, 2003).

Even though predatory microarthropods did not appear to reduce prey density, the species composition did change through time which may be linked to either direct predation pressure and/or intra/inter-specific interactions among prey. Either one of these may have led to preferred prey species no longer surviving and certain species dominating the system toward the end, causing a decrease in predator abundance. As this change in species composition over time occurred in both the control and high density treatment coupled with the fact that gamasid mites are thought to be generalist feeders (Karg, 1993; Beaulieu and Weeks, 2007) the latter might be a more likely explanation of the decrease in predator density. The change in species composition over time (due to intra-specific interactions among prey) to smaller-bodied species (results not shown), may have made them inaccessible to the larger predators by occupying smaller pore spaces. Of course, the decrease in predator pressure might have had a knock-on effect on fast reproducing, species becoming more abundant thus compensating for increased predation. Although an increased relative abundance of smaller sized species in response to environmental change is not uncommon (Lindo, 2015), an explanation as to why a shift in community composition occurred in this particular instance can only be speculative.

A change through time in some feeding groups which occur in both the control and high density treatment (D8), of a decrease in herbofungivores (Symphypleona juveniles), and an increase in decomposer species also occurred. The decrease in Symphypleona juveniles may simply be a reduction in food such as plant derived tissue and algae over time (Chahartaghi et al., 2005). Collembola of the genus *Folsomia* and *Tomocerus,* which compose the primary decomposer feeding group, exemplify the difficulty in assigning a taxon to any one feeding group when, for example, *Folsomia* is documented not only as feeding on detritus but also to be fungivorous and nematophagous (in the case of *Folsomia candida*) (Lee and Widden, 1996; Chahartaghi et al., 2005; Fountain and Hopkin, 2005). This species (*F. candida*) not alone makes use of a multitude of food sources but is known to be parthenogenetic and to do well in microcosms (Usher and Stoneman, 1977) which may account for the observed increase in abundance over time.

As various components such as microbial biomass C were affected by compression, a look at compressed samples alone (not shown) show a similar pattern/trend as discussed above. This is interesting as it suggests that a system with reduced food and abundance of enchytraeids (not shown) as well as some shifts in proportions of feeding groups, can lead to the same overall density. The fact that a reduction in volume of soil pore space does not affect overall microarthropod abundance supports the idea that something other than space as a resource might be limiting at this particular scale. That said, studies on the effect of compaction on various aspects of the soil generally suggest that compaction results in a reduction in density and diversity due to the reduction in habitable pore space (Heisler and Kaiser, 1995; Battigelli et al., 2004). In this study, although compression reduced the volume of various sized pores typically inhabited by microarthropods, it did not significantly affect their overall abundance. The accessibility and habitability of soil pores (for Collembola at least) is determined by a number of physical aspects of the soil including volume of habitable pore space and connectivity of these pores (Hopkin, 1997). As the latter was not measured in this study, the results can only be partially explained, if at all. Given the few explanatory variables to work with, it is worth noting that in this study the microcosms were undisturbed intact soil cores taken directly from the field and microarthropods entered the soil cores from the surface after compression i.e. fauna were not exposed to mechanical stress (with the exception of the controls). Therefore, unlike many other studies investigating the effects of compaction (which generally restructure and/or modify the soil) the presence of roots and natural cracks may have resulted in higher connectivity between soil pores, potentially undoing the effect of compression. In addition, the influence of microarthropods on soil structure may generally be underrepresented in the literature, as is discussed in the few studies that have explored their influence on soil aggregation (Maaß et al. 2015, Siddiky et al. 2012).

Although compression did not significantly affect total microarthropod abundance, in certain instances community composition were affected. For example, as herbofungivores were solely composed of Symphypleona juveniles it is likely that their abundances were reduced as they tend to be epiedaphic (probably residing in and among the few remaining grass shoots on the surface of the soil core/microcosm) (Hopkin, 1997). The Collembola of the genus *Tomocerus* on the other hand are relatively large (up to 4.5 – 6.2 mm) which may account for the reduction in decomposers due to compression. In fact, when we look at rough estimates of Collembola biomass (not shown) in conjunction with species composition we can see that similar numbers of organisms of smaller sized species appear to dominate at the end of the experiment. This supports aboveground terrestrial and aquatic studies which suggest that smaller individuals may be more abundant than larger ones, corresponding to the amount of available habitat (Kampichler 1995, White *et al*. 2007).

## 5 Conclusions

## 6 Acknowledgements

We would like to thank Coimbra University, Portugal and Teagasc Centre Johnstown Castle, Ireland for help with the PLFA analysis. We would also like to thank Matthias Bacher, Teagasc Centre, Johnstown Castle, Ireland for help with the analysis of soil porosity. Thank you to the land owner Thomas Wall for letting us access and take soil samples from his field. Special thanks to Phil Fanning and Cormac Mc Conigley who helped collect the soil cores.

This research did not receive any specific grant from funding agencies in the public, commercial, or not-for-profit sectors.

## Notes

### Competing Interest Statement

The authors have declared no competing interest.

